# Sampling and ranking of protein conformations using machine learning techniques do not improve the quality of rigid protein-protein docking

**DOI:** 10.1101/2025.05.13.652389

**Authors:** Roman Kyrylenko, Ihor Koleiev, Illia Savchenko, Taras Voitsitskyi, Roman Stratiichuk, Vladyslav Husak, Semen Yesylevskyy, Serhii Starosyla, Alan Nafiiev

## Abstract

Rigid docking remains the most popular method of predicting protein-protein interactions in cases when experimental 3D structures of the complexes are not available. The docking often relies on known unbound (Apo) protein structures, which may differ significantly from their bound (Holo) forms. Modern machine learning (ML) based conformational sampling techniques allow generating ensembles of functionally relevant protein structures, which may be closer to their Holo forms and thus could improve the outcomes of the classical rigid protein-protein docking. Here, we sampled conformations of the protein subunits in 30 complexes from the novel PINDER dataset with two state-of-the-art ML-based techniques and evaluated their docking performance using several physics-based, data-based, and ML-based scoring functions. We showed that such conformational sampling rarely produces structures that are closer to the Holo conformations than the corresponding Apo ones. Moreover, even when such conformations are generated, none of the tested scoring functions were able to prioritize and rank them correctly. Our work highlights critical limitations in the current ML-enhanced rigid protein-protein docking workflows and emphasizes the need for new approaches that can better utilize the potential of modern techniques for conformational generation and scoring.

## 1. Introduction

Protein–protein interactions (PPIs) are fundamental to nearly all cellular functions and molecular causes of diseases (1,2). Understanding the three-dimensional structure of the protein-protein interfaces is especially crucial for structure-based drug discovery workflows (3). However, experimental structural characterization of the PPIs remains challenging and costly. Consequently, computational methods, particularly *in silico* protein-protein docking, play an increasingly important role in predicting the structure of protein complexes (4–6).

Rigid protein-protein docking still remains a method of choice in most cases when experimentally determined bound (Holo) structures of the protein complexes are unavailable. In such cases, docking must rely on available unbound (Apo) conformations or computationally predicted monomer structures (e.g., from AlphaFold2 (7) or custom homology modeling). These structures, however, may differ significantly from the biologically relevant conformation adopted upon binding due to induced fit or conformational selection mechanisms (8) (Fig. 1).

**Figure 1.**
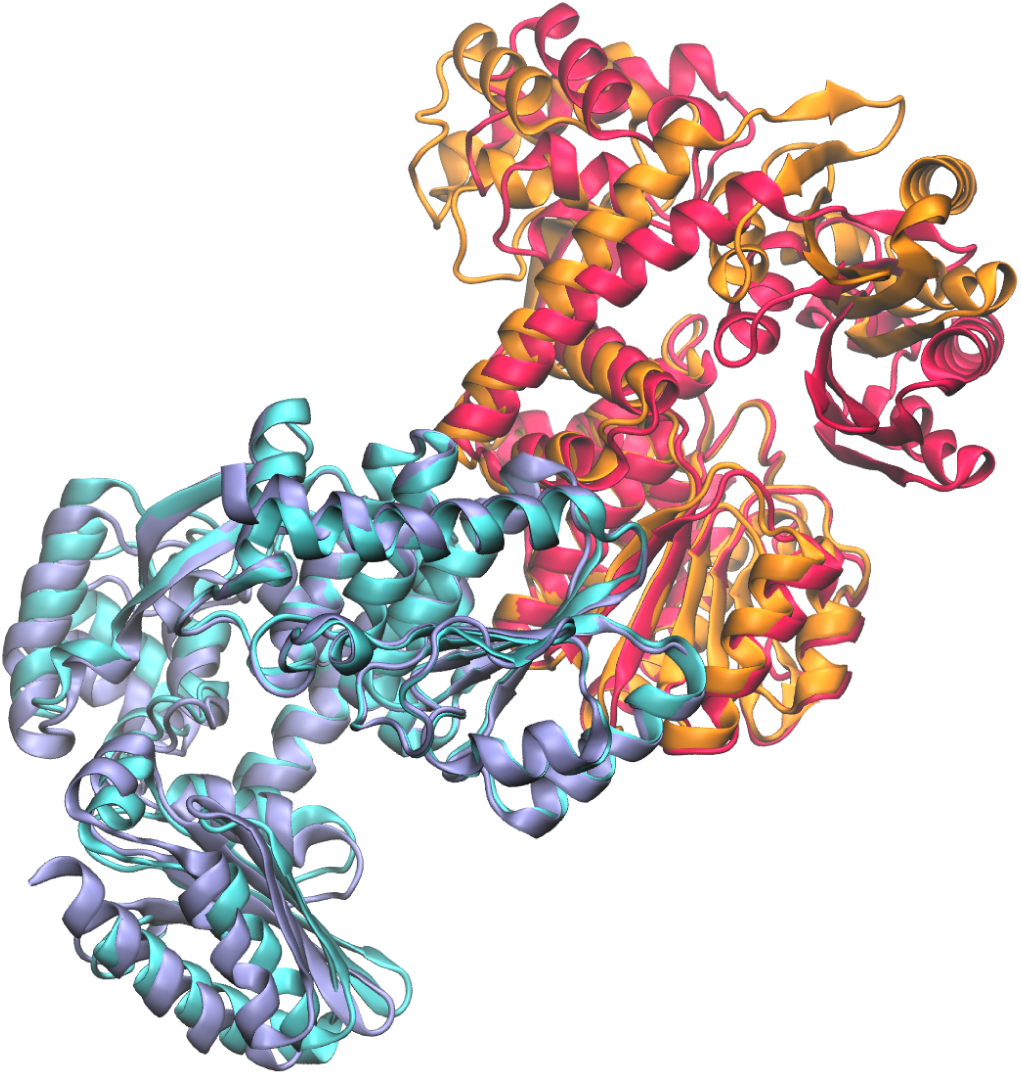
An example of the large conformational change occurring upon the formation of the PPI in the Single-stranded DNA exonuclease homodimer complex (UniProt ID of the apo subunits - Q58241, PDB ID of the holo complex - 7YKV) included in the PINDER-AF2 dataset. Docked apo subunits are shown in red and blue, subunits in the crystallized holo complex are orange and cyan, respectively.

**Figure 2.**
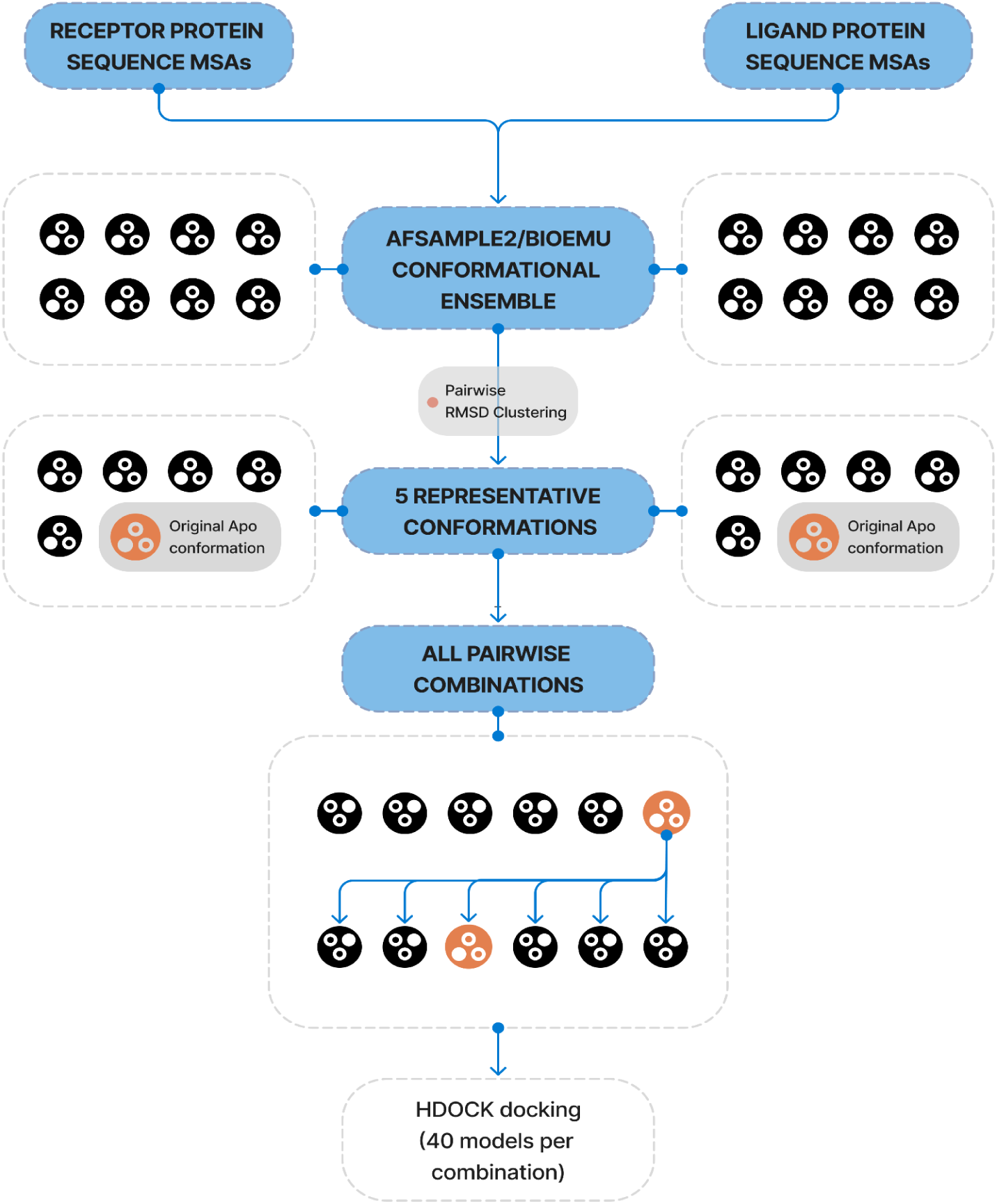
General scheme of the pipeline with consecutive sampling, clustering, and docking stages.

The classical ways to sample protein conformations beyond the rigid Apo structures include such approaches as Molecular Dynamics (MD) and Monte Carlo (MC) simulations usually paired with enhanced sampling techniques, such as Replica Exchange, Metadynamics, Umbrella sampling, etc. (10). Despite being quite robust, these methods are computationally demanding and suffer from the convergence issues. Recent advances in machine learning (ML) offer a promising alternative for efficiently sampling protein conformational ensembles (11–14). These methods have the potential to explore protein conformational space faster than classical approaches and are shown to contain functionally relevant conformations.

The most widely adopted approach, which is at the core of currently state-of-the-art methods in this field, is based on the hidden potential of the AlphaFold (AF) model to predict different possible conformations of the target protein by introducing random or systematic variations on different stages of its inference algorithm. This includes modifying input MSA in various ways (AFCluster (14), MSAsubsample (15), AFTraj (13)); introducing noise into AF inference (AFSample (12), AFSample2 (11), AFTraj); or not using template information or its part during the inference (AFSample, AFSample2, AFTraj).

It is logical to hypothesize that ML-generated conformational ensembles may contain near-Holo conformations, thereby providing better starting points for docking and potentially bridging the performance gap currently observed when using only Apo structures. To test this hypothesis, we decided to use AFSample2 (11) and BioEmu (16) ML sampling techniques. AFSample2 is a trusted technique that clearly outperforms other related methods based on MSA variations, while BioEmu is one of the latest developments in the field, tailored for maximal performance and thus discussed and explored extensively in the community.

This study investigates whether combining ML-based conformational sampling with a classical, efficient rigid-body docking algorithm (HDOCK (17)) can improve the prediction of protein complexes. We employed two distinct ML-based sampling methods (AFSample2 and BioEmu) to generate conformational ensembles for proteins from the most advanced and challenging PPI benchmark dataset to date - PINDER-AF2 Apo (18). These ensembles were then used in docking simulations.

However, generating potentially improved conformations is not enough. It is of critical importance to correctly identify these near-native conformations within a large ensemble of docked structures, which presents a major scoring challenge. Standard docking scoring functions, often developed or calibrated using experimental Holo structures or static docked models, may not be accurate enough at distinguishing correct interfaces from the vast number of non-native decoys generated during the ensemble docking. This may create a ‘scoring bottleneck’ when the PPI prediction fails not because of the absence of correctly docked conformations, but because of their incorrect scoring and prioritization.

Given this uncertainty, we evaluated the ability of diverse scoring strategies to identify successful predictions within the docking results. This included the HDOCK intrinsic score alongside representative physics-based, knowledge-based, and ML-based rescoring functions (17,19–22). Testing this panel was crucial because it remains unclear *a priori* which, if any, current scoring approach can successfully prioritize near-native complexes derived from ML-sampled Apo structures. Docking success was rigorously assessed using criteria established in CAPRI (15,16).

Our results indicate that while ML sampling occasionally generated conformations structurally closer to the Holo state than the input Apo structure, this rarely translated into improved rigid-body docking success regardless of the used scoring function. Critically, even when superior conformations were sampled and led to accurate docked complexes (as revealed by oracle scoring), none of the tested scoring functions prioritized them correctly. In this work, we highlight these concurrent limitations in both state-of-the-art ML conformational sampling and PPI scoring. We discuss the need for new approaches, potentially integrating flexibility more directly or developing ensemble-aware scoring functions, to better harness structural diversity for accurate PPI prediction.

## 2. Methods

### 2.1. Protein conformation sampling

#### 2.1.1. AFSample2 protocol

As in AF, the sampling of conformations with AFSample2 includes the MSA-building stage and the structure prediction stage. For the MSA building, the default AFSample2 procedure was used (which is basically a default AF procedure) - i.e., the Jackhmmer (23), HHblits (24), and HHsearch (25) are used for MSA construction from default AF databases (with the full version of the BFD database).

For the structure prediction, we used the following configuration: not using templates for structure prediction, the maximum number of recycles is 3, enabling the dropout for the inference, the fraction of random MSA masking is 0.2, using weights of 5 AF’s monomer models (which results in 5 predicted conformations per 1 iteration of structure prediction), the number of structure prediction iterations is 10 (plus 30 additional iterations later). The AFSample2 produces structures with all heavy atoms, so no sidechain reconstruction is needed. For structure minimization, we used the OpenMM (26) molecular dynamics suite, which handled both protonation and energy minimization. For minimization, we constrained the positions of the backbone heavy atoms and performed 500 steps of minimization.

#### 2.1.2. BioEmu protocol

BioEmu also depends on the MSA information, however, it uses ColabFold’s (27) MSA generation procedure, which is based on MMseqs2 (28) and utilizes the open MSA server, which stores the sequence databases. For BioEmu’s inference, we used the default ColabFold’s MSA generation procedure and the original version of model weights (it should be emphasized that BioEmu could undergo updates since the time of writing, and new configurable parameters could appear).

The structures sampled with BioEmu contain only backbone heavy atoms, so the side chains were further reconstructed using BioEmu’s side chain reconstruction procedure, which is based on H-Packer (29). The procedure also includes subsequent protonation of structures with the PropKa (30) and minimization procedure, which is based on OpenMM. As both the H-Packer-based side chain reconstruction and the minimization take significant time per sample (∼20 seconds), we first performed the clustering of the conformations (see the details below) and then reconstructed and minimized only the cluster centers.

### 2.2. Protein conformations clustering

To systematically categorize protein conformations sampled by AFSample2 or BioEmu, we applied Hierarchical agglomerative clustering using pairwise root mean square deviation (RMSD) as the structural dissimilarity metric. This approach enables the identification of representative conformations that preserve the diversity of the sampled conformational ensemble (31).

Initially, the RMSD distance matrix for all sampled protein conformations was computed. Clustering was then performed using the agglomerative hierarchical method with complete linkage. In this context, "complete linkage" ensures that the maximum distance between pairs of points in different clusters is minimized, promoting the formation of compact and well-separated clusters.

To identify representative conformations for each cluster, we computed the average RMSD of each member structure to all other structures within the same cluster. The structure with the minimal average RMSD was selected as the representative cluster center.

Manipulations with protein structures and RMSD computations were performed with the custom scripts based on the MolAR molecular modeling library (32).

### 2.3. Protein-Protein docking

Docking was conducted using HDOCK (17), a standalone, Fast Fourier Transform(FFT)-based protein-protein rigid docking program that efficiently explores the rotational and translational space. HDOCKlite, derived from the comprehensive HDOCK framework, was selected for its high computational efficiency, robust prediction accuracy, and open-source accessibility, making it particularly well-suited for large-scale docking of ML–sampled protein conformations.

In addition to its well-established performance in CAPRI (33–36), HDOCK was recently benchmarked on the novel PINDER dataset, a curated and challenging collection of protein–protein interactions used to assess docking accuracy in diverse scenarios. In this evaluation, HDOCK demonstrated strong performance, outperforming or matching the accuracy of several leading docking algorithms, further validating its utility for structure-based interaction modeling in realistic and nontrivial cases.

Docking was performed in template-free mode with default scoring parameters. The overall docking pipeline involved sampling alternative conformations for each protein, selecting structurally diverse cluster representatives, and docking all-vs-all combinations of these representatives, including Apo forms.

### 2.4. Scoring functions

Following the generation of docking complexes using HDOCK, we applied a rescoring strategy aimed at prioritizing the most relevant structures. Our setup required a scoring scheme capable of handling diverse conformational states and capturing subtle interface differences that may not be evident in rigid or single-conformer docking.

To address this challenge, we independently rescored each docked structure using the following scoring functions:

● **HDOCK Scoring**: Used as the baseline, this knowledge-based scoring function combines empirical energy terms, residue contact potentials, and shape complementarity. While tightly integrated with the docking engine, it often favors shape-based metrics and may overlook conformational flexibility (17).
● **PRODIGY-Cryst**: This scoring method does not predict binding affinity per se, but instead discriminates between biologically relevant interfaces and crystallographic artifacts. It uses features derived from interfacial residue contacts and crystal structure statistics, making it especially valuable for identifying likely physiological complexes from rigid docking results (37).
● **dMaSIF (Deep Mesh-based Surface Interaction Fingerprinting)**: A deep learning–based scoring method that computes molecular surfaces directly from atomic coordinates and uses geometric convolutions to identify interaction-relevant surface features. dMaSIF efficiently captures local interface properties, possibly making it well-suited for distinguishing near-native from non-native complexes in rigid docking scenarios (22).
● **PIsToN (Protein Binding Interfaces with Transformer Networks)**: PIsToN leverages structure-aware transformer architecture to encode both sequence and 3D spatial context. PIsToN also uses additional empirical energy terms, calculated by the FireDock (38) refinement module, as inputs to the neural network to construct a hybrid machine learning model (21).
● **HADDOCK3 Score**: A physics-based scoring function derived from the HADDOCK3 suite that evaluates models based on a weighted sum of van der Waals, electrostatics, desolvation energy, and buried surface area (19,20).

### 2.5. Evaluation dataset

To evaluate the performance of our ensemble-based docking pipeline, we employed the PINDER-AF2 Apo benchmark dataset, a part of the broader PINDER dataset. Unlike older PPI datasets that often suffer from data leakage between training and test sets due to reliance on sequence similarity clustering, PINDER utilizes interface residue–based structural clustering, which substantially reduces the risk of overlap and information leakage (18). This ensures a more realistic and generalizable assessment of docking and scoring methods. By focusing on interface-level structural similarity rather than global sequence identity, PINDER provides a rigorous benchmark for assessing the true generalization capability of ML-based methods.

The PINDER-AF2 test set consists of 180 protein–protein complexes, all of which were deposited after the AlphaFold-Multimer training cutoff, ensuring no training data contamination when evaluating AlphaFold2-derived or AlphaFold2-compatible methods (39). From this dataset, we selected 30 complexes with 32 unique proteins for which experimentally determined Apo structures are available, allowing us to simulate docking conditions using unbound conformations (Apo-docking) in addition to disassembled complexes (Holo-docking).

### 2.6. Evaluation metrics

To assess the quality of docking predictions and the effectiveness of different scoring functions within our ensemble docking pipeline, we employed a Top-N success rates evaluation strategy based on DockQ scoring.

DockQ is a widely adopted continuous quality metric that integrates multiple CAPRI criteria — including interface RMSD, ligand RMSD, and the fraction of native contacts — into a single score ranging from 0 (poor) to 1 (perfect). DockQ scores were used to assign CAPRI-like quality categories (Incorrect, Acceptable, Medium, High) to individual docked decoys, enabling interpretable and standardized comparisons across methods(40,41).

Similarly to the existing PINDER benchmark, we evaluated Top-N performance, reporting the percentage of complexes for which at least one decoy of a given CAPRI quality was found among the Top-1, Top-5, or Top-40 ranked models, based on each scoring function. An additional “Oracle” metric was also reported, representing the upper limit of success for each method by checking whether any decoy of acceptable or better quality was present across the full candidate pool, regardless of ranking. This metric estimates the performance of a hypothetical perfect scoring function.

## 3. Results

### 3.1. Characterisation of the PINDER-AF2 subset

To assess the potential influence of the source structures (Holo vs Apo) on docking performance, we began by examining 30 complexes selected from the PINDER-AF2 benchmark for which Apo protein structures are available.

#### 3.1.1 Sequence Length Differences

As a preliminary check, we analyzed sequence length differences between the Holo and Apo forms for both receptor and ligand subunits. Minor discrepancies can arise due to construct truncations, alternate splicing, or unresolved residues in crystal structures. The histogram in Supplementary Figure 3 shows the distribution of sequence length differences (Holo - Apo) across the subunits of 30 tested complexes. Overall, the majority of cases cluster around zero, with only a few outliers showing length differences exceeding ±20 residues. This suggests that Holo and Apo forms in this subset are mostly consistent in terms of sequence coverage and that significant global structural rearrangements are unlikely due to truncation alone.

#### 3.1.2 HDOCK Score Differences

To assess how docking performance varied between structural forms, we docked each complex twice using either Holo or Apo structures of the subunits. We then computed the HDOCK score difference (Holo - Apo) for each complex, shown in Supplementary Figure 1. For the 30 complexes used, the HDOCK score varied from -185 to -983 for Holo-Holo and -186 to -638 for Apo-Apo. As the HDOCK score is an approximation of binding energy, lower scores indicate better predicted binding.

The score difference distribution exhibits a modest left skew, with a mean difference of −75.45, indicating that Holo–Holo docking configurations were generally scored more favorably. While this preference is not uniform across all complexes, the lower quartile values (Q10 = −456.41, Q25 = −192.44) reveal substantially better scores when using the Holo structures. Corresponding descriptive statistics are summarized in Supplementary Table 1.

#### 3.1.3 PRODIGY-Cryst Score Differences

To evaluate the biological relevance of the obtained interfaces, we also rescored each docked complex using PRODIGY-Cryst, which estimates the likelihood of an interface being biologically meaningful (vs. crystallographic artifact). Score differences (Holo - Apo) across the 30 complexes are shown in Supplementary Figure 2. The distribution again shows a slight leftward trend, with PRODIGY-Cryst assigning higher biological relevance to Holo interfaces in many cases. These values support the interpretation that Holo-based models are often perceived as more biologically plausible, although the effect is subtle and varies across complexes (Supplementary Table 1).

### 3.2. Protein conformation sampling and clustering

The conformations for all unique monomers (n=36) found in the PINDER-AF dataset were generated using either AFSample2 or BioEmu. For the homodimer complexes, where Apo-structures contained different numbers of resolved amino acids, we considered each monomer sequence as unique. We generated 50 samples with AFSample2 and 500 samples with BioEmu.

The quality of generated samples differs significantly between the methods. For AFSample2, no artifacts were encountered, while for BioEmu, multiple issues were observed. For a significant fraction of the structures, BioEmu’s built-in OpenMM-based minimization procedure failed, even though the side-chain reconstruction was successful. We traced the problem to incompatibility between PropKa-protonated structures and MDTraj, which is used for processing between the side chain reconstruction and minimization, and fails to correctly assign bonds for non-standard residues. Also, we encountered problems with the missing second carboxylic oxygen on the C-terminus. To fix these issues, the PDBFixer (the OpenMM helper script) was used to add the missing atoms and to make protonation compatible with OpenMM.

We also noticed chain ruptures in some conformations, which turned out to be the effect of abnormally large lengths of the peptide bonds. After inspecting the minimization procedure used by BioEmu, we found out that the positions of backbone heavy atoms are restrained, which doesn’t allow OpenMM minimization to fix such bonds. We removed these restraints from the minimization procedure, which resolved the issue.

To obtain a structurally diverse subset of conformations for docking, we selected five representative structures per protein using agglomerative hierarchical clustering as described in Methods. To assess the structural variability of resulting ensembles, we computed the distribution of pairwise RMSD values among selected structures (Figure 3).

**Figure 3.**
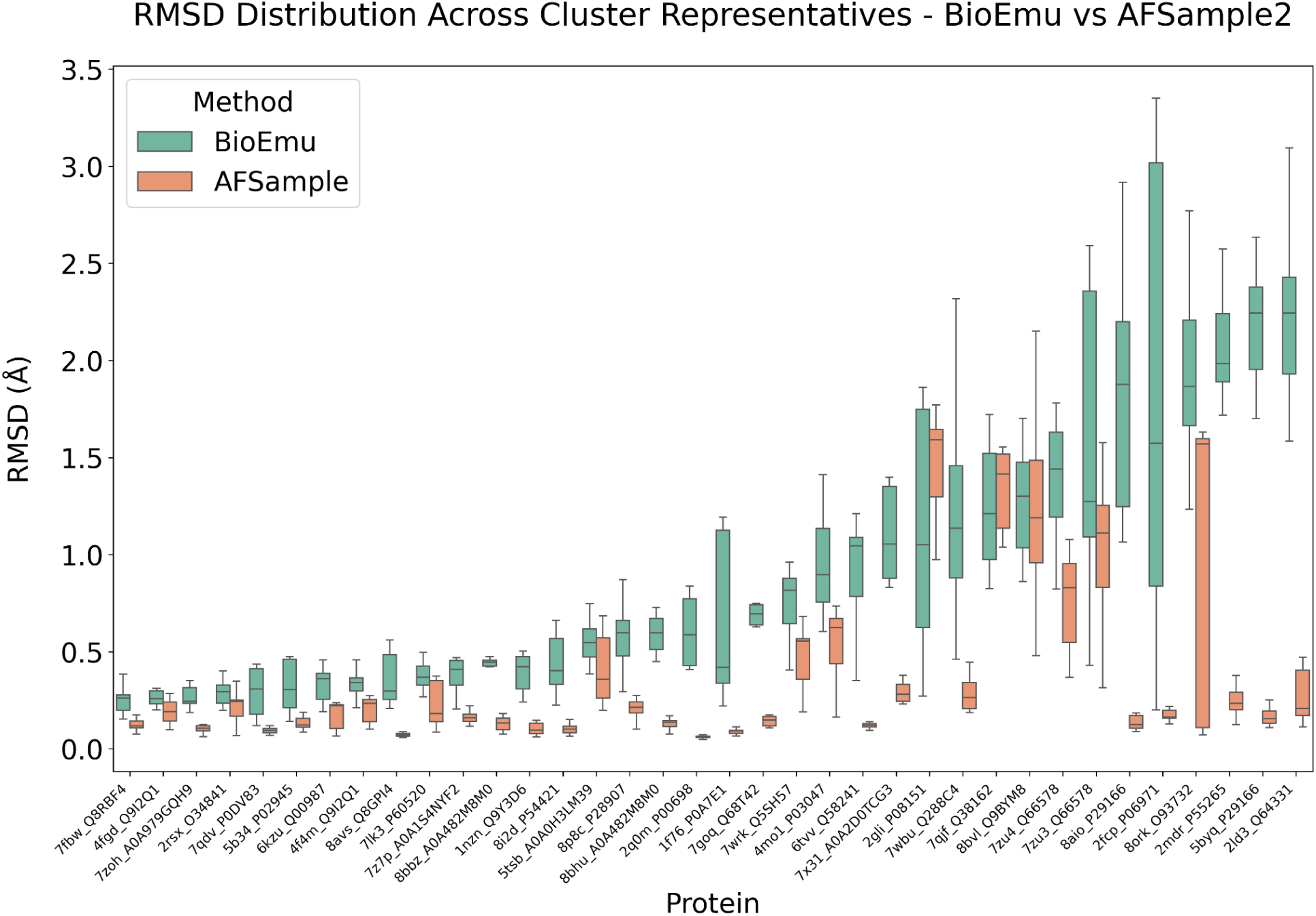
RMSD distribution among selected representative conformations generated by BioEmu and AFSample2 for all studied proteins (identified by complex PDB ID and UniProt ID).

The results show that BioEmu produces broader RMSD distributions, with higher medians and maximum pairwise deviations in most cases, indicating greater structural diversity among selected conformations. In contrast, AFSample2 forms more compact sets of conformations, with narrower RMSD ranges and lower median values. This difference may be partially explained by the larger number of generated conformations (500 for BioEmu vs 50 for AFSample2). Although BioEmu may offer broader sampling coverage, it also introduces a wider range of structural variability, which can include both useful diversity and potentially irrelevant noise.

### 3.3. Docking of predicted structures

#### 3.3.1. Influence of conformational sampling on docking accuracy

The original Apo structures were added to five representative generated conformations of each protein subunit. All 36 combinations per complex were docked using HDOCK. 40 models per run were generated and evaluated by the protocol from the original PINDER study. This yielded a total of 43,200 docking predictions across 30 benchmark complexes.

Table 1 summarizes the docking success rates based on DockQ CAPRI classification, comparing AFSample2 and BioEmu-generated conformations against established Apo-Apo docking baselines. Results were assessed at Top-1, Top-5, and Top-40 predictions. An “Oracle” column captures the maximum DockQ score obtainable per complex, assuming perfect ranking of the 1,440 models per complex.

**Table 1.**
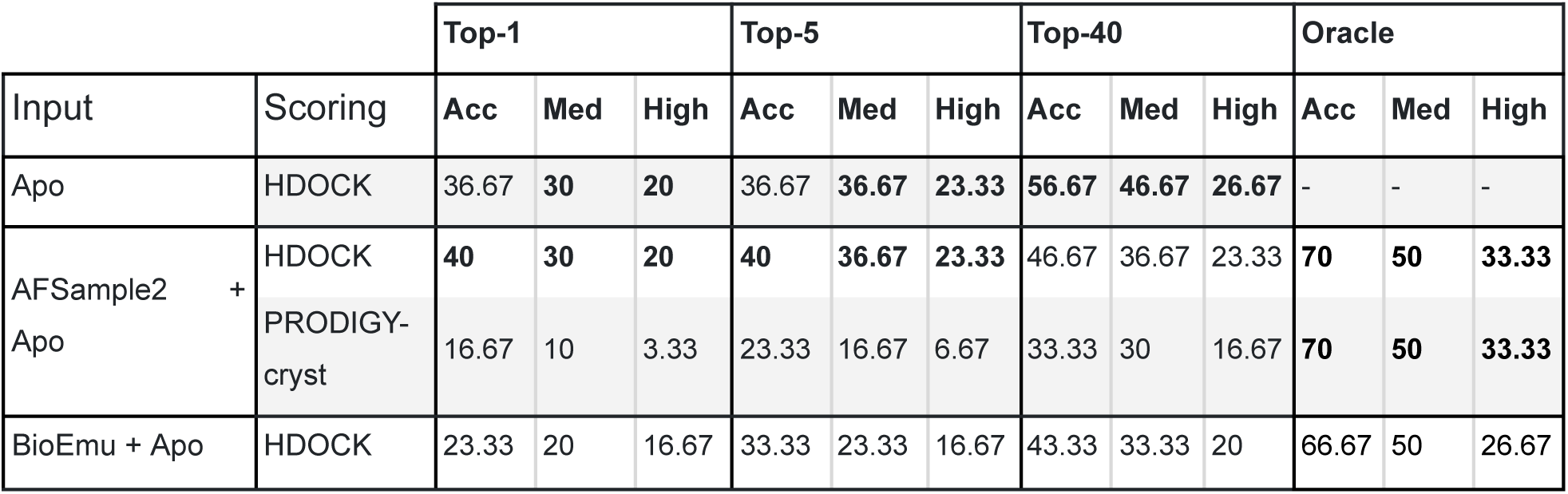
CAPRI DockQ classification across different docking pipelines using Apo PINDER-AF2 benchmark (N=30). Each method evaluated Top-1, Top-5, and Top-40 predictions. Results are reported as percentages of targets with Acceptable (Acc), Medium (Med), or High-quality predictions in terms of DockQ CAPRI classification. The Oracle column represents the best-case scenario (highest DockQ per complex over all models generated by docking). The best values for the respective columns are marked in bold.

| Input | Scoring | Top-1 |  |  | Top-5 |  |  | Top-40 |  |  | Oracle |  |  |
| --- | --- | --- | --- | --- | --- | --- | --- | --- | --- | --- | --- | --- | --- |
|  |  | Acc | Med | High | Acc | Med | High | Acc | Med | High | Acc | Med | High |
| Apo | HDock | 36.67 | **30** | **20** | 36.67 | **36.67** | **23.33** | **56.67** | **46.67** | **26.67** | - | - | - |
| AFsample2 + Apo | HDock | **40** | **30** | **20** | **40** | **36.67** | **23.33** | 46.67 | 36.67 | 23.33 | **70** | **50** | **33.33** |
|  | PRODIGY-cryst | 16.67 | 10 | 3.33 | 23.33 | 16.67 | 6.67 | 33.33 | 30 | 16.67 | **70** | **50** | **33.33** |
| BioEmu + Apo | HDock | 23.33 | 20 | 16.67 | 33.33 | 23.33 | 16.67 | 43.33 | 33.33 | 20 | 66.67 | 50 | 26.67 |

It is evident that AFSample2 sampling provided a marginal improvement in the docking accuracy. Top-1 and Top-5 improved from 36.67% to 40%, which corresponds to an improvement in only one of the docked complexes. The Top-40 performance was lower than the original Apo-only docking, indicating that the relevant conformations were often displaced by less suitable models, reducing the overall quality. BioEmu performed even worse with lower Top-1, Top-5, and Top-40 metrics compared to both Apo-only and AFSample2 runs. This suggests that the structural diversity generated by BioEmu may be less compatible with rigid-body docking, or that it introduces conformations that are structurally less favorable under HDOCK’s scoring scheme.

However, the Oracle values for generated complexes are much higher than for apo structures, which means that both AFSample2 and BioEmu generate a significant proportion of high-quality complexes. This discrepancy underscores a key limitation: HDOCK’s default scoring function is unable to effectively rank the best docked structures in generated conformational ensembles.

To explore alternative scoring strategies, we evaluated the same set of structures using PRODIGY-Cryst. While PRODIGY-Cryst was able to differentiate between Apo and Holo complexes, its utility as a scoring function appeared to be limited. The Top-1 and Top-5 success rates with PRODIGY-based ranking were noticeably worse than those with HDOCK, and the Top-40 were comparable. These findings highlight that despite its biological relevance, PRODIGY-Cryst is not optimized for evaluating rigid-body docking ensembles.

It is possible to conclude that conformational sampling with both AFsample2 and BioEmu produces protein conformations that may improve the rigid docking quality, however, neither the default HDOCK scoring function nor interface-focused metrics like PRODIGY-Cryst are effective in correctly ranking these conformations.

#### 3.3.2. Analysis of sampled conformations

To assess whether ML conformational sampling can better approximate the bound (Holo) state of a protein–protein interface, we performed a detailed analysis of generated structures. We defined the binding site in the original Holo complex as all Cα atoms within 10 Å of the other partner. Using this residue selection, we aligned the corresponding regions of the Apo structure and the ML-sampled conformations onto the Holo subunit and computed the root mean square deviation (RMSD) of all heavy atoms within the interface. This enabled a direct comparison of how closely sampled conformations resemble the Holo PPI compared to the Apo conformation. The results are shown in Figures 4 and 5.

**Figure 4.**
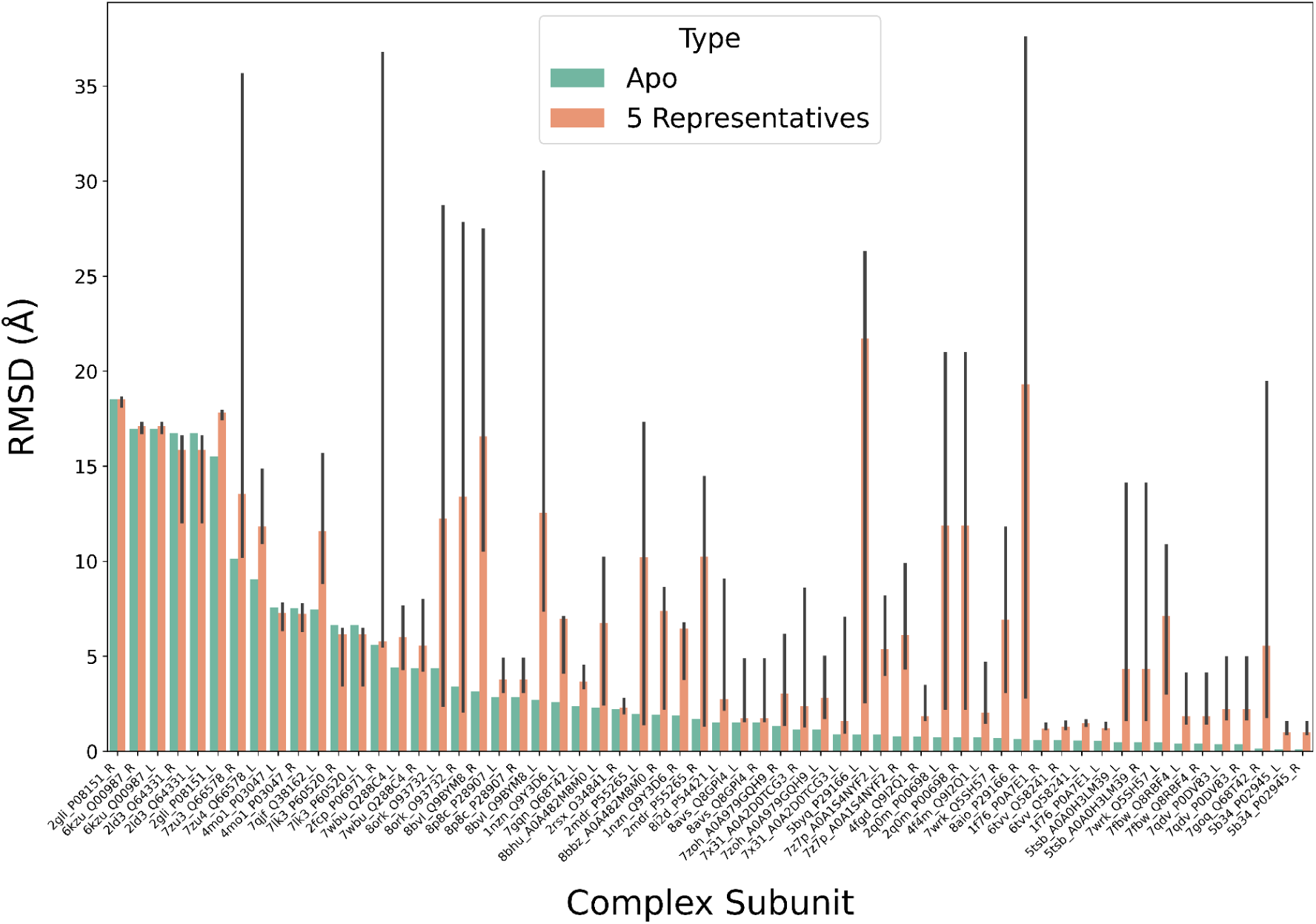
RMSD values computed for the binding interface residues between the Holo structure and either Apo or 5 representative BioEmu-sampled conformations for all studied complex subunits. Green bars represent RMSD with the Apo structure. Red bars represent RMSD with the cluster centroids. The bars show the median RMSD, while the whiskers show the minimum and maximum values. Binding interfaces were defined as the residues within 10 Å of the other partner in the target Holo complex. Subunits are identified by the PDB ID of the complex, the UniProt ID of a protein, and whether it is in the Receptor or Ligand position.

**Figure 5.**
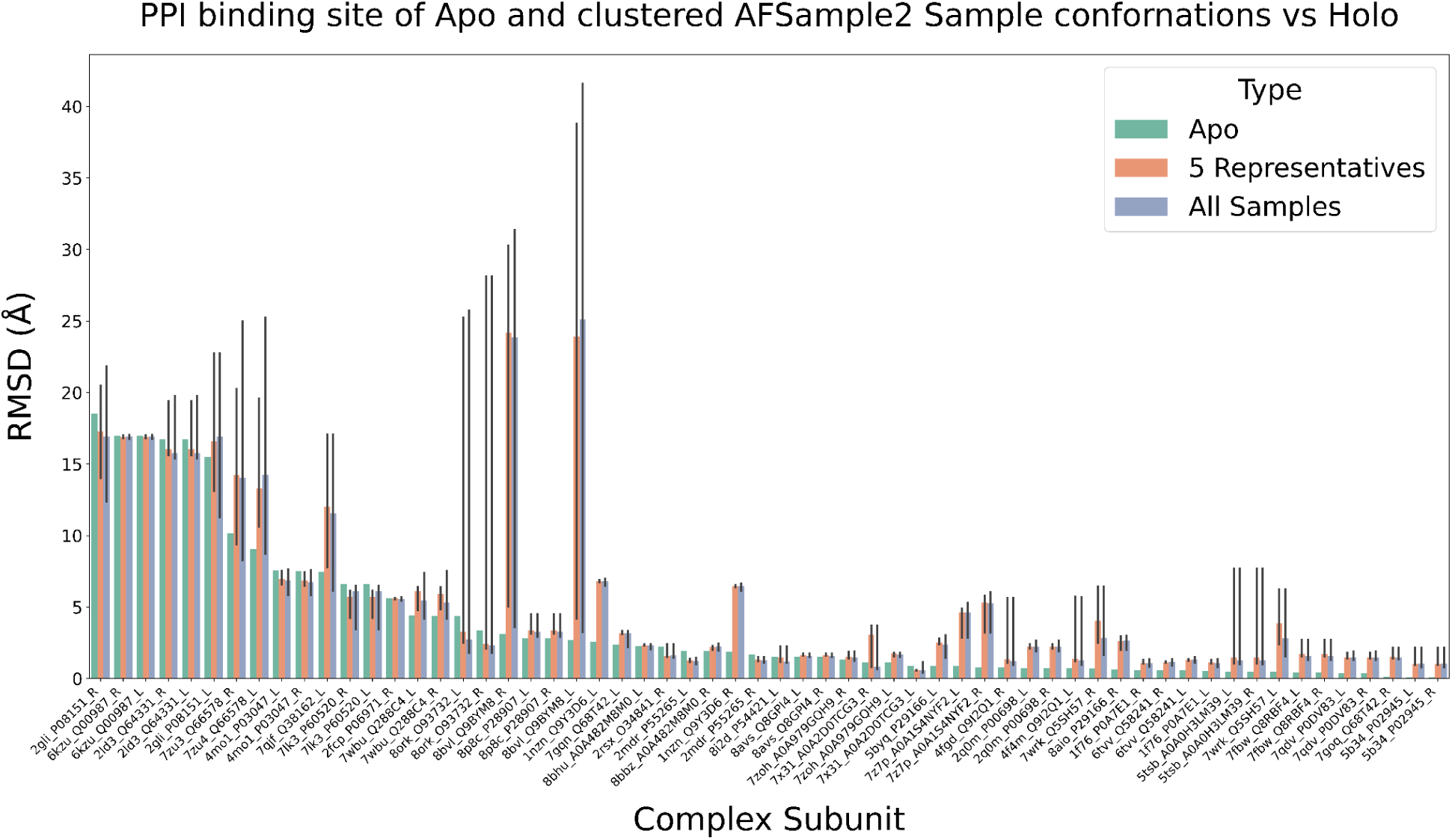
RMSD values computed for the binding interface residues between the Holo structure and Apo, 5 representative, or all 50 AFSample2-sampled conformations for all studied complex subunits. Green bars represent RMSD with the Apo structure. Red bars represent RMSD with the cluster centroids. Blue bars represent RMSD with all sampled conformations. The bars show the median RMSD, while the whiskers show the minimum and maximum values. Binding interfaces were defined as the residues within 10 Å of the other partner in the target Holo complex. Subunits are identified by the PDB ID of the complex, the UniProt ID of a protein, and whether it is in the Receptor or Ligand position.

It is clearly visible that the majority of sampled conformations, regardless of the method, are not closer to the Holo conformation than the original Apo structure. In many cases, conformational sampling produces structures that diverge from the Holo structure much more than the original Apo structure. This is especially pronounced in the BioEmu dataset, where nearly all sampled conformations are farther from the Holo interface than the Apo form. This suggests that BioEmu may not effectively explore functionally relevant conformations in the context of PPIs, which could also explain the poor performance of BioEmu’s sampled conformations in the rigid docking (Table 1).

In contrast, the AFSample2 demonstrates much better RMSD distribution, with some sampled structures approaching or even slightly improving upon the Apo conformation in 26 of 60 subunits. However, these improvements are still marginal, and sampled conformations still do not fully capture structural features of the bound state. We also noted that the RMSD distribution of five representative structures closely resembles the distribution over all 50 samples, suggesting that clustering successfully captures the conformational diversity present in the sampled ensemble.

Our findings emphasize the challenges of sampling functionally relevant conformations of PPIs using current structure prediction ML methods.

#### 3.3.3. Detailed analysis of HDOCK scoring function performance

To further investigate the influence of conformational sampling on docking performance, we conducted a detailed analysis of docking for 30 PINDER-AF2-Apo protein-protein complexes. For each complex, three different approaches were used:

● Apo - models consisting of the original unbound (Apo) conformations of both subunits.
● Cluster - models consisting of subunits sampled with AFSample2, where conformations were selected based on clustering, with no prior knowledge of the target binding site.
● BestConf - models consisting of AFSample2-generated conformations, but the selection was guided by identifying structures with binding sites closest to the Holo conformation in terms of RMSD. These represent ideal selections from available sampled conformations.

We compared DockQ scores and HDOCK energy scores of the docked complexes to evaluate the quality of HDOCK scoring. For each complex, we computed the difference in DockQ and HDOCK score between the best sampled model (either Cluster–Cluster or BestConf–BestConf) and the best baseline Apo model, selected by DockQ. This allowed us to assess not only whether conformational sampling improved the quality of predicted structure but also whether such improvements were captured by the scoring function.

As shown in Table 2, the Cluster approach is superior to Apo in 14 of 30 cases, indicating a closer approximation to the native complex by clustered structures. However, these improvements were not reflected in the HDOCK energy scores. In most cases, a better structural model, as judged by DockQ, received a worse or comparable HDOCK score to the baseline structure. This suggests that HDOCK’s scoring function does not consistently reward geometrically accurate complexes generated by conformational sampling. This decoupling between structural accuracy and scoring highlights a critical limitation of HDOCK: even when sampling successfully generates high-quality models, they may not be selected correctly.

**Table 2.**
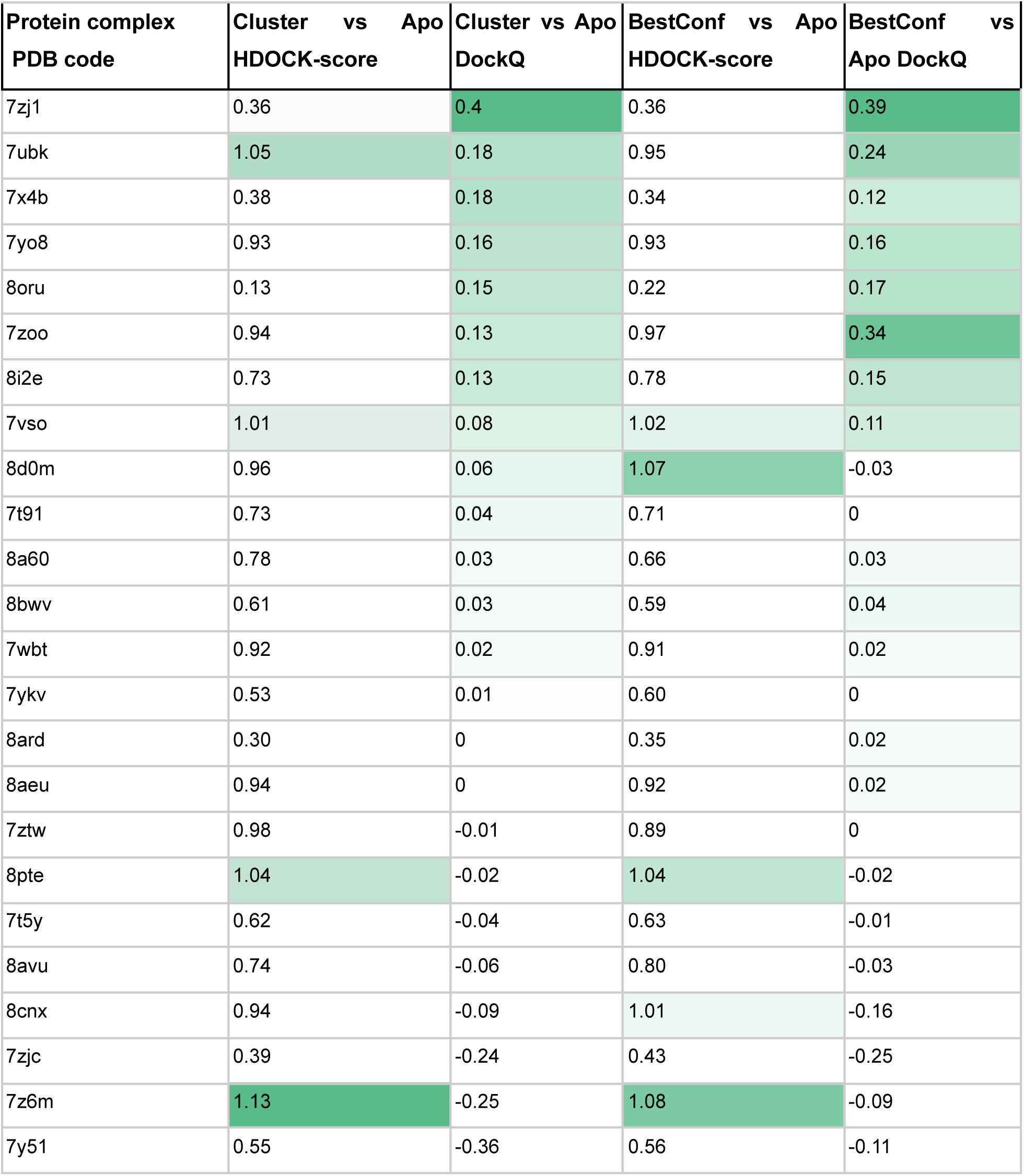
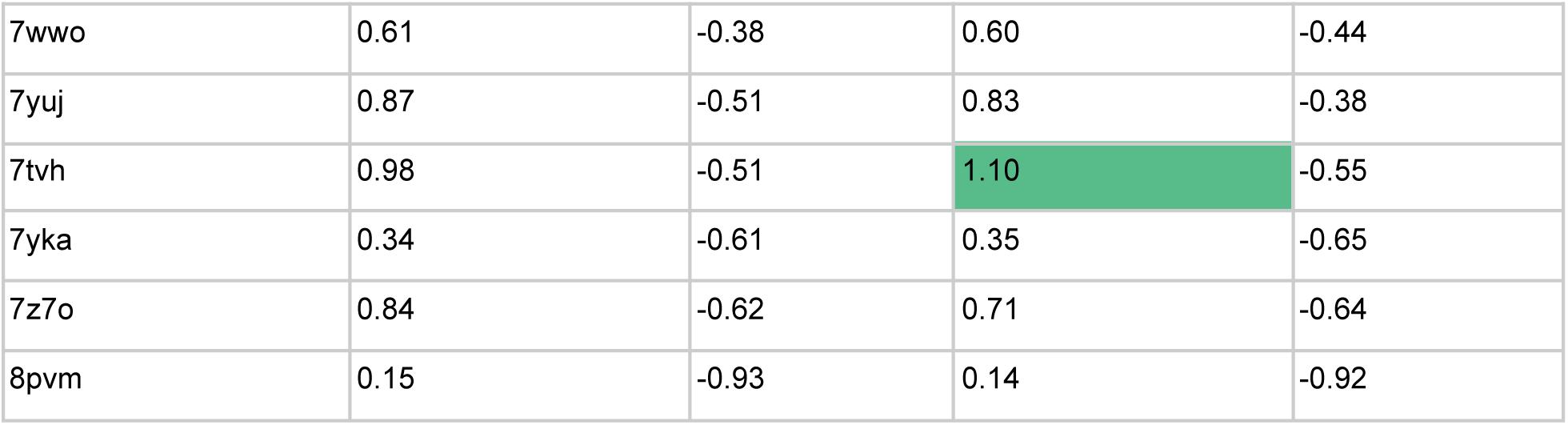
Comparison of docking performance across the studied complexes using three modeling strategies: Apo (Apo-Apo complexes, where both subunits are in their original unbound conformations), Cluster (Cluster-Cluster complexes, where both subunits are sampled via AFSample2 and selected through clustering), and BestConf (BestConf-BestConf complexes, where subunits are selected from sampled ensembles based on binding-site RMSD to the Holo reference). The table shows the differences in DockQ and HDOCK score between Cluster and Apo, and between Best and Apo. DockQ values represent the absolute difference (Other DockQ - Apo DockQ), while HDOCK values are reported as relative ratios (Other HDOCK / Apo HDOCK), where a value of 1 indicates no change. Green shading marks cases where Cluster or BestConf complexes outperform Apo.

It should also be noted that the selection of representative conformations based solely on their similarity to holo conformations in the binding interface (BestConf) did not always lead to improvements in DockQ scores, suggesting that better binding site conformation does not always correlate with better overall conformation quality.

These findings emphasize the need for improved scoring strategies or complementary ranking approaches that can detect and prioritize structurally correct sampled conformations more reliably.

#### 3.3.4. Exploring alternative scoring functions

In an effort to address the limitations observed with the original HDOCK scoring function, we explored alternative scoring methods. Our initial hypothesis was that the HDOCK scoring function might inherently favor the Apo states, which typically represent more energetically stable and native-like contact distributions compared to ML-sampled structures. This prompted us to investigate other scoring methods that could potentially provide a more balanced assessment across both Apo and predicted conformations.

We followed an approach of Shirali et al. (42), where a range of scoring functions from different classes — physics-based, knowledge-based, empirical, and machine learning-based — were evaluated on eight benchmark PPI datasets. Their findings highlighted the strong performance of machine learning-based methods, dMaSIF (22) and PIsToN (21), which consistently outperformed traditional scoring approaches across diverse benchmarks. Based on these results, we selected dMaSIF and PIsToN as representative ML-based methods, and complemented them with HADDOCK3 as a physics-based alternative. The results obtained with these scoring models are presented in Table 3.

**Table 3.** Comparing scoring performance across multiple methods using the PINDER-AF2-Apo benchmark (N=30) based on DockQ CAPRI classification. Scored with various strategies. Docking performed with HDOCK. Results are reported as percentages of targets with Acceptable (Acc), Medium (Med), or High-quality predictions in terms of DockQ CAPRI classification. The best values for the respective columns are marked in bold.

| Input | Scoring | Top-1 |  |  | Top-5 |  |  | Top-40 |  |  | Oracle |  |  |
| --- | --- | --- | --- | --- | --- | --- | --- | --- | --- | --- | --- | --- | --- |
|  |  | Acc | Med | High | Acc | Med | High | Acc | Med | High | Acc | Med | High |
| Apo | HDock | 36.67 | **30** | **20** | 36.67 | **36.67** | **23.33** | **56.67** | **46.67** | **26.67** | - | - | - |
| AFSample2 | HDock | **40** | **30** | **20** | **40** | **36.67** | **23.33** | 46.67 | 36.67 | 23.33 | **70** | **50** | **33.33** |
|  | PlsToN | 13.33 | 6.67 | 0 | 30 | 20 | 10 | 43.33 | 36.67 | 20 | **70** | **50** | **33.33** |
|  | dMaSiF | 6.67 | 6.67 | 3.33 | 16.67 | 6.67 | 3.33 | 36.67 | 20 | 6.67 | **70** | **50** | **33.33** |
|  | HADDOCK3-EM | 20 | 16.67 | 13.33 | 26.67 | 26.67 | 16.67 | 46.67 | 36.67 | 20 | **70** | **50** | **33.33** |
|  | HADDOCK3-MD | 10 | 10 | 6.67 | 23.33 | 20 | 6.67 | 43.33 | 30 | 16.67 | **70** | **50** | **33.33** |

None of the tested techniques managed to match or surpass the performance of the original HDOCK score. This discrepancy was especially pronounced in the Top-1 and Top-5 ranked complexes, where HDOCK consistently performed better. To our surprise, dMaSIF and PIsToN showed the worst overall performance, which is most likely caused by the characteristics of the used dataset. The PINDER-AF2 subset represents a temporal cutoff that excludes all data before the training of AlphaFold-Multimer. However, both dMaSIF and PIsToN were trained prior to this cutoff. This eliminates the possibility of data leakage and shows that the generalizability of these scoring functions to new, unseen protein complexes is poor.

For the HADDOCK3, two variations suggested by the documentation were tested: HADDOCK3-EM (scoring after energy minimization) and HADDOCK3-MD (scoring after short restrained molecular dynamics in explicit solvent). While HADDOCK3-EM did show improvement over the ML-based approaches, it still lagged behind the original HDOCK score. Surprisingly, the HADDOCK3-MD variant performed worse than the HADDOCK3-EM variant. The reason for this is not entirely clear. One possible explanation is that MD simulations drive the structures away quickly, but they are too short to sample the return to correct conformations.

These results highlight an unexpected robustness of the HDOCK scoring function within our specific docking pipeline and surprisingly bad performance of both the state-of-the-art ML scoring functions and sophisticated physics-based techniques on the PINDER-AF2 dataset.

#### 3.3.5. Evaluation of extended conformational sampling

To further explore the limitations of conformational sampling, we conducted an experiment in which we extended the number of conformations generated by AFSample2. Specifically, we increased the number of sampled conformations per subunit from 50 to 200, aiming to enhance conformational diversity and increase the likelihood of finding Holo-like structures. The expectation was that by enlarging the ensemble, we could enrich the sampling space with more functionally relevant conformations, potentially leading to improved docking outcomes. However, the results did not support this hypothesis. As summarized in Table 4, performance metrics across Top-1, Top-5, and Top-40 DockQ classifications actually decreased with the larger ensemble, compared to the baseline sampling (50 conformations per subunit).

**Table 4.**
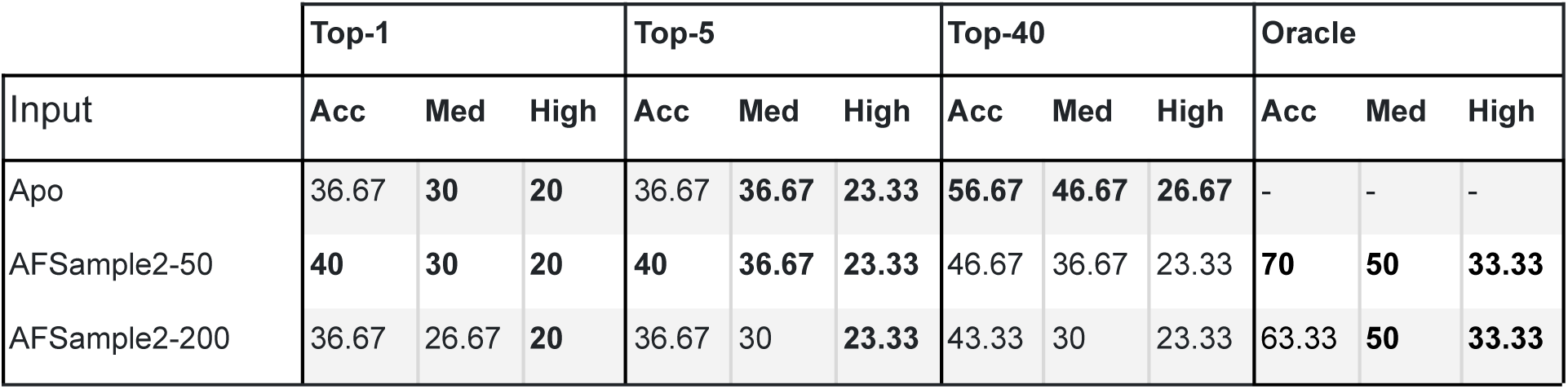
HDOCK performance with additional AFSample2 sampled conformations. AFSample2-N denotes N generated conformations per subunit, which are then clustered to get 5 representative conformations. Results are reported as percentages of targets with Acceptable (Acc), Medium (Med), or High-quality predictions in terms of DockQ CAPRI classification. The best values for the respective columns are marked in bold.

|  | Top-1 |  |  | Top-5 |  |  | Top-40 |  |  | Oracle |  |  |
| --- | --- | --- | --- | --- | --- | --- | --- | --- | --- | --- | --- | --- |
| Input | Acc | Med | High | Acc | Med | High | Acc | Med | High | Acc | Med | High |
| Apo | 36.67 | **30** | **20** | 36.67 | **36.67** | **23.33** | **56.67** | **46.67** | **26.67** | - | - | - |
| AFSample2-50 | **40** | **30** | **20** | **40** | **36.67** | **23.33** | 46.67 | 36.67 | 23.33 | **70** | **50** | **33.33** |
| AFSample2-200 | 36.67 | 26.67 | **20** | 36.67 | 30 | **23.33** | 43.33 | 30 | 23.33 | 63.33 | **50** | **33.33** |

Furthermore, the maximum achievable performance — the Oracle score assuming perfect ranking — did not improve. This indicates that additional conformations introduce more noise and dilute the ensemble without any consistent advantage. These results suggest that increasing the number of sampled conformations alone is insufficient to improve docking success. Without more effective strategies for selecting and ranking near-native structures, a larger sampling space may only increase the difficulty of identifying useful structures. This finding further underscores the limitations of current sampling methods and highlights the need for structure-aware clustering or selection methods to extract meaningful diversity while avoiding redundancy and irrelevant outliers.

### 3.4. Limitations

While this study provides a detailed analysis of conformational sampling and scoring strategies for rigid protein–protein docking, several limitations should be acknowledged. First, the size of the benchmark dataset - 30 complexes from the PINDER-AF2-Apo subset - limits the statistical power of some of our findings. Although the dataset was carefully curated to ensure temporal and structural independence from the training sets of evaluated ML techniques, a broader and more diverse dataset would help to generalize the observed trends.

Second, the dataset is heavily skewed toward homodimeric interactions, with 28 out of 30 complexes being homodimers. This overrepresentation may bias conclusions about sampling and scoring performance, particularly since homodimers often tend to present more regular and less flexible interfaces than heterodimeric complexes (43). Future work should evaluate these approaches on more heterogeneous PPIs.

Third, while HDOCK was used as the primary docking engine due to its favorable CAPRI performance and accessibility, other state-of-the-art docking frameworks, particularly flexible docking methods, were not tested. Including additional docking algorithms could reveal whether the scoring limitations we observed are specific to HDOCK or generalize to all protein-protein docking methods.

Finally, although basic agglomerative clustering was used to select representative sampled conformations, this approach may not fully capture the structural diversity or relevance of the ensemble. More advanced or task-specific clustering strategies, such as interface-aware, energy-weighted, or ML-guided clustering, could better identify conformations that are close to the Holo complexes. Exploring and benchmarking a wider range of clustering and selection methods remains an important direction for improving the effectiveness of ensemble-based docking pipelines.

## 4. Discussion

This work presents a comprehensive assessment of ML-based conformational sampling and different scoring strategies for improving the results of rigid protein–protein docking. Using the PINDER-AF2-Apo benchmark, we tested whether generated protein conformations can enhance docking performance beyond that achievable with experimentally derived Apo structures.

Our results demonstrate that, despite the theoretical potential of ML-based sampling methods, such as AFSample2 and BioEmu, current implementations do not consistently produce protein conformations that outperform Apo structures when used for docking. In many cases, sampled conformations deviate more from the Holo binding interface than the initial Apo structures. For BioEmu, the generated conformations were typically inferior to the Apo baselines, while for AFSample2, they were on par but not significantly better than the original unbound structures. Even in the cases where improved interface geometries were generated, they were not properly selected by the docking scoring functions, regardless of their nature (physics-based, data-based, or ML-based).

A central finding of this study is that the docking scoring function itself, not the structures of docked proteins, plays a critical role in limiting protein-protein docking accuracy. We show that the HDOCK score, which is widely used and performs relatively well in the rigid-body docking benchmarks, fails to consistently recognize near-native complexes derived from ML-generated conformations of subunits. While HDOCK score performed best among the other tested scoring methods, it seems to be biased toward the Apo structures over predicted alternatives. This is supported by our detailed analysis of scoring behavior across conformational categories (Apo–Apo, Cluster–Cluster, and BestConf–BestConf), which revealed that structurally superior models (as judged by DockQ) often received worse HDOCK scores and were therefore excluded from top selections.

To determine whether alternative scoring approaches could overcome this limitation, we benchmarked several modern scoring functions representing different paradigms: dMaSIF and PIsToN (machine learning–based), and HADDOCK3-EM and HADDOCK3-MD (physics-based). While claimed to be highly promising in the previous studies, these methods performed poorly on our benchmark. ML-based scoring functions demonstrated an inability to generalize over the new PINDER-AF2-Apo dataset, which is distinct from their training datasets with a minimized possibility of data leakage. Physics-based HADDOCK3 scoring showed some improvement over the ML-based methods, but still fell short of HDOCK in ranking performance. Oracle scores indicate that high-quality models exist in the sampled space but are not effectively identified by any of the tested scoring techniques.

## 5. Conclusion

Our findings reveal two core challenges in using modern ML conformational sampling and ranking techniques with the rigid protein-protein docking. First, ML conformational sampling methods struggle to generate conformations of the subunits, which resemble correct Holo structures in the complex. Second, even if such conformations are obtained, the scoring functions—regardless of whether they are knowledge-based, physics-based, or ML-based—struggle to recognize and prioritize them correctly. This hinders the discovery of near-native structures in large, diverse conformational ensembles and limits the practical applicability of docking workflows based on the structures predicted by ML techniques. Future research should prioritize the development of scoring functions that are robust to conformational variability, trained on broader distributions of modern predicted structures, and capable of generalizing beyond legacy datasets.

## 6. Author contributions

AN served as the conceptual lead and originator of the core idea. RK and IK carried out literature research, protein-protein docking, wrote the scripts, performed clustering, scoring, and comprehensive result analysis, including visualization. TV supervised the project, participated in the design of the study, provided technical support, and designed and developed a protein-protein docking pipeline. IS implemented and optimized the AFSample2 and BioEmu conformational sampling pipelines and contributed to the manuscript editing. RS and VH analyzed the dataset and provided general support. SY coordinated the work, critically assessed the results, and guided the analysis. SS and AN designed the study and controlled its progress. The manuscript was written by RK and SY.

## 8. Supporting Information

Additional details, figures, and statistical analysis of the Results (DOCX).

## 8. Acknowledgements

S.Y. has received funding through the grant MSMT-355/2025-16 from the Ministry of Education, Youth and Sport of the Czech Republic and through the MSCA4Ukraine project 101101923, which is funded by the European Union. Views and opinions expressed are, however, those of the author(s) only and do not necessarily reflect those of the European Union. Neither the European Union nor the MSCA4Ukraine Consortium as a whole nor any individual member institutions of the MSCA4Ukraine Consortium can be held responsible for them.

The authors acknowledge the use of the large language model Gemini 2.5 Pro for proofreading and stylistic editing of the manuscript.

