## Supplementary Information for "Sampling and ranking of protein conformations using machine learning techniques do not improve the quality of rigid protein-protein docking"

*Roman Kyrylenko* [1*], *Ihor Koleiev* [1,2], *Illia Savchenko* [1], *Taras Voitsitskyi* [1,2], *Roman Stratiichuk* [1,5], *Vladyslav Husak* [1,6], *Semen Yesylevskyy* [1,2,3,4], *Serhii Starosyla* [1], *Alan Nafiiev* [1].

[1] Receptor.AI Inc., 20-22 Wenlock Road, London N1 7GU, United Kingdom.

[2] Department of Physics of Biological Systems, Institute of Physics of The National Academy of Sciences of Ukraine, 46 Nauky Ave., 03038, Kyiv, Ukraine.

[3] Institute of Organic Chemistry and Biochemistry, Czech Academy of Sciences, CZ-166 10 Prague 6, Czech Republic.

[4] Department of Physical Chemistry, Faculty of Science, Palacký University Olomouc, 17. listopadu 12, 771 46 Olomouc, Czech Republic.

[5] Department of Biophysics and Medical Informatics, Educational and Scientific Centre “Institute of Biology and Medicine”, Taras Shevchenko Kyiv National University, 64 Volodymyrska Str., 01601, Kyiv, Ukraine.

[6] Department of Cellular, Computational and Integrative Biology, The University of Trento, Via Sommarive 9, 38123 Povo (Trento), Italy

*

HDOCK score distributions

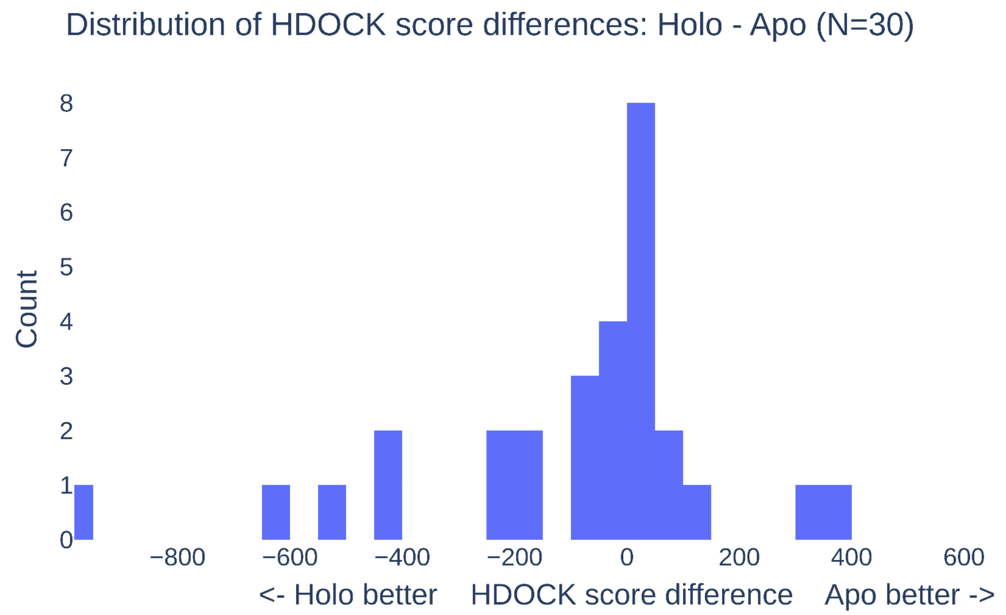


**Supplementary Figure 1**. Distribution of the HDOCK score differences (Holo - Apo) across the studied complexes.

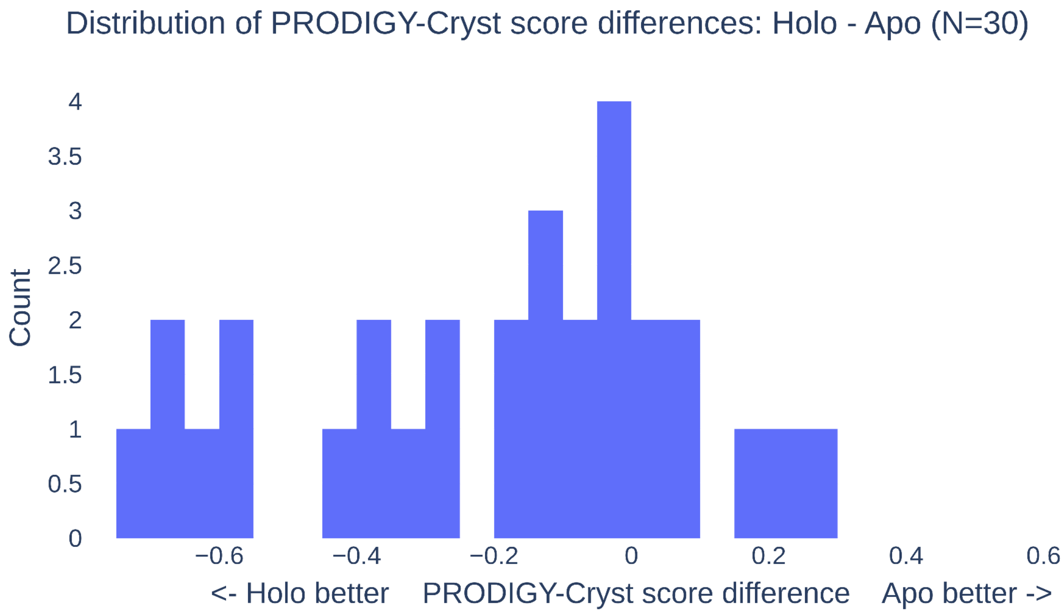


**Supplementary Figure 2**. Distribution of the PRODIGY-Cryst score differences (Holo - Apo) across the studied complexes. Negative values imply higher confidence for Holo-based predictions.

### Details of the protein conformational sampling

#### AFSample2

The time complexity of AFSample is comparable to that of AlphaFold and depends primarily on sequence length and MSA depth. For all but one sequence in the PINDER-AF2-Apo dataset, MSA generation (on a 16 vCPU, 64 GB RAM CPU machine) and structure prediction (on a GPU machine with an NVIDIA T4, 8 vCPUs, and 30 GB RAM) required approximately 15 and 51 hours, respectively. For the longest sequence (899 amino acids), MSA generation exceeded the 64 GB RAM limit and was completed using a 128 GB RAM machine in ~2 hours; structure prediction on the same GPU setup took ~15 hours.

#### BioEmu

In BioEmu, the batch_size_100 parameter defines the batch size normalized to a sequence length of 100 residues and is used internally to determine the actual batch size during inference. We evaluated different batch_size_100 values on a GPU with an NVIDIA T4 (16 GB VRAM). At values of 20 and 100, GPU utilization remained at 100%, while memory usage increased from 2 GB to 10 GB, respectively. However, prediction time at batch_size_100 = 100 (~90 seconds) was approximately five times longer than at 20 (~18 seconds), indicating that increased memory usage does not yield performance gains and that GPU compute throughput is the limiting factor. We set batch_size_100 to 100 and generated 500 conformations per protein. The total runtime for the full dataset was approximately 44 hours—demonstrating a key advantage of BioEmu over AFSample2: an approximately 20-fold reduction in runtime.

### Details of the characterisation of the PINDER-AF2 subset

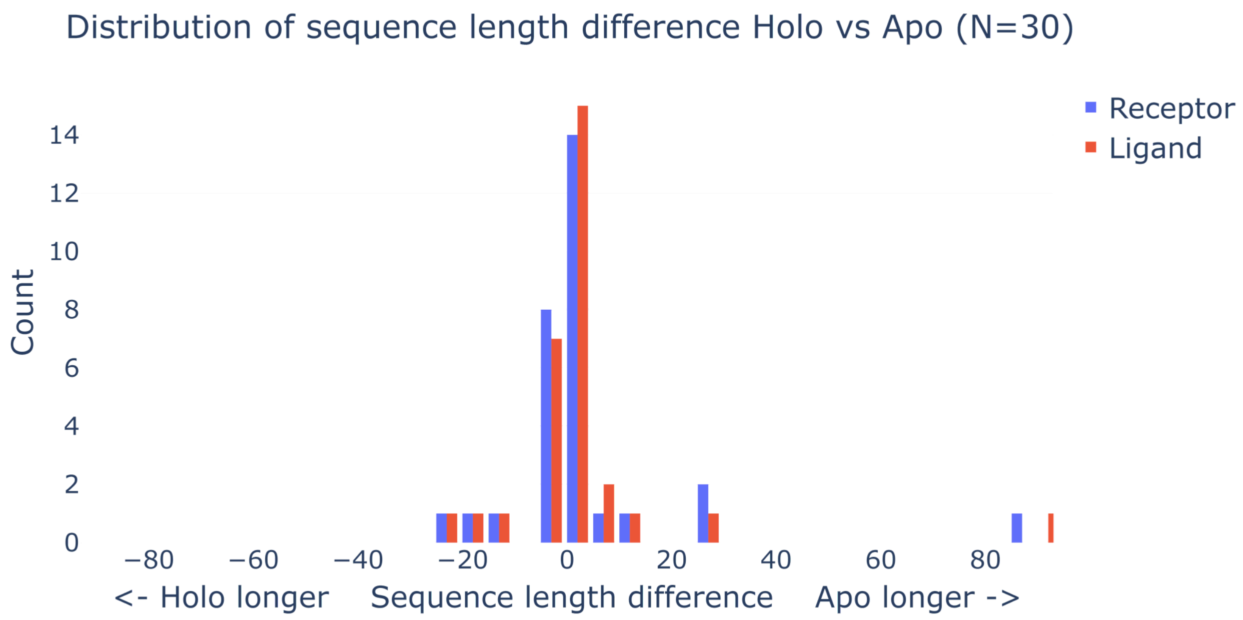

**Supplementary Figure 3**. Distribution of sequence length differences (Holo - Apo) for receptors (larger subunits of complexes) and ligands (smaller subunits of complexes) across the studied PINDER-AF2 complexes.

To assess the robustness of the observed trends between Holo and Apo docking performance, we applied two non-parametric statistical tests to the distribution of score differences (Holo – Apo) across the 30 complexes:

- The Wilcoxon signed-rank test was used to evaluate whether the median of the paired score differences significantly deviated from zero. This test is appropriate for paired, non-normally distributed data.
- The Sign test was used to determine whether one structural form (Holo or Apo) consistently outperformed the other by simply counting the number of positive vs. negative score differences. The sign test evaluates whether the observed imbalance is unlikely to have occurred by chance.

Statistical testing indicated that HDOCK score Holo structures preference trend is not robustly significant. A Wilcoxon signed-rank test yielded a p-value of 0.114 which is above the standard threshold for significance. A sign test showed that only 16 out of 30 complexes had negative differences ($p = 0.428$), reinforcing the conclusion that while a directional preference exists, it lacks statistical support under conventional thresholds (Supplementary Table 2).

Unlike HDOCK, the PRODIGY-cryst score leftward skew trend was statistically significant. The Wilcoxon signed-rank test returned a p-value of 0.0005. The sign test also confirmed significance ($p = 0.0026$). These results indicate that PRODIGY-Cryst systematically evaluates Holo–Holo docking models as more biologically plausible than Apo–Apo models. (Supplementary Table 2)

**Supplementary Table 1**. Summary of score differences between Holo–Holo and Apo–Apo docking models across 30 PINDER-AF2 complexes (<0 - Holo scored better, >0 - Apo scored better).

| Statistic | HDOCK Score Difference | PRODIGY-Cryst Score Difference |
|---:|---:|---:|
| **Mean** | −75.45 | −0.20 |
| **Q10** | −456.41 | −0.64 |
| **Q25** | −192.44 | −0.39 |
| **Q50 (Median)** | −5.99 | −0.11 |
| **Q75** | 18.33 | −0.01 |
| **Q90** | 139.16 | 0.08 |

**Supplementary Table 2**. Statistical evaluation of score differences (Holo – Apo) for HDOCK and PRODIGY-Cryst scores across 30 PPI complexes.

| Test Type | HDOCK Score Difference | PRODIGY-Cryst Score Difference |
|---:|---:|---:|
| **Wilcoxon signed-rank** | p = 0.114 (not significant) | p = 0.0005 (significant) |
| **Sign test (n negative)** | 16/30 → p = 0.428 (ns) | 23/30 → p = 0.0026 (significant) |
